# Hyperprolactinemia in a male pituitary androgen receptor knockout mouse model is associated with a female-like pattern of lactotroph development

**DOI:** 10.1101/2020.08.05.236927

**Authors:** Laura O’Hara, Helen C. Christian, Paul Le Tissier, Lee B. Smith

## Abstract

Circulating prolactin concentration in rodents and humans is sexually dimorphic. Estrogens are a well-characterised stimulator of prolactin release. Circulating prolactin fluctuates throughout the menstrual/estrous cycle of females in response to estrogen levels, but remains continually low in males. We have previously identified androgens as an inhibitor of prolactin release through characterisation of males of a mouse line with a conditional pituitary androgen receptor knockout (PARKO) which have an increase in circulating prolactin, but unchanged lactotroph number. In the present study we aimed to specify the cell type that androgens act on to repress prolactin release. We examined lactotroph-specific, Pit1 lineage-specific and neural-specific conditional AR knockouts, however they did not duplicate the high circulating prolactin seen in the pituitary androgen receptor knockout line, suggesting that the site of androgen repression of prolactin production was another cell type. Using electron microscopy to examine ultrastructure we showed that pituitary androgen receptor knockout male mice develop lactotrophs that resemble those seen in female mice, and that this is likely to contribute to the increase in circulating prolactin. When castrated, pituitary androgen receptor knockout males have significantly reduced circulating prolactin compared to intact males, which suggests that removal of circulating estrogens as well as androgens reduces the stimulation of pituitary prolactin release. However, when expression of selected estrogen-regulated anterior pituitary genes were examined there were no differences in expression level between controls and knockouts. Further investigation is needed into prolactin regulation by changes in androgen-estrogen balance, which has implications not only in the normal sexual dimorphism of physiology but also in diseases such as hyperprolactinemia.

## Introduction

Prolactin is a hormone primarily associated with the process of lactation. However, since its initial discovery it has been found to be involved in over 300 different physiological processes in both males and females (Bole-Feysot et al., 1998). It is theorised that prolactin may act as a general ‘pregnancy and lactation’ hormone, promoting processes in the body necessary to aid successful production and feeding of offspring (Grattan and Le Tissier, 2015). Prolactin is produced by the lactotroph cells of the anterior pituitary. Control of lactotroph prolactin production and release appropriate for the organism’s physiological state is achieved from a balance of stimulation and inhibition. The main prolactin inhibitory factor is dopamine, produced by the tuberoinfundibular dopaminergic (TIDA) neurones of the hypothalamus which make contact with blood vessels in the median eminence and release dopamine to be transported in the hypophyseal portal circulation to the anterior pituitary (Ben-Jonathan and Hnasko, 2001, Freeman et al., 2000). Since lactotrophs spontaneously release prolactin when they are isolated *in vitro*, or when the pituitary gland is isolated from the hypothalamus *in vivo*, it is accepted that prolactin release is under tonic inhibition by dopamine (Grattan and Le Tissier, 2015). Prolactin releasing factors of hypothalamic origin include oxytocin, thyrotrophin-releasing hormone and vasoactive intestinal peptide (VIP) (Freeman et al., 2000).

Estrogen is also a well-characterised stimulator of prolactin release. Circulating prolactin concentration is sexually dimorphic. Prolactin release fluctuates throughout the estrous cycle of rodents and the menstrual cycle of humans. In rats, circulating prolactin peaks in the afternoon of proestrus as estrogen levels are rising before the LH surge (Hawkins et al., 1975). In humans, prolactin is significantly higher during the ovulatory and luteal phases than during the follicular phase (Franchimont et al., 1976). Prolactin release in male rats does not cycle, and is similar to that of the lowest concentration in cycling females at diestrus (Amenomori et al., 1970).

We have previously identified androgens as an inhibitor of prolactin release through characterisation of a mouse line with ablation of androgen receptor in the pituitary (pituitary androgen receptor knockout: ‘PARKO’) (O’Hara et al., 2015). Male PARKO mice have an increase in both pituitary *Prl* transcript and circulating prolactin concentration, despite no increase in lactotroph number. Androgen receptor is present in 71% of gonadotrophs, 50% of lactotrophs, 45% of thyrotrophs, 25% of corticotrophs and 16% of somatotrophs (O’Hara et al., 2015). The PARKO model, driven by Foxg1-Cre, is an ablation of androgen receptor in all of the cells of the pituitary gland. Since lactotrophs produce prolactin and most express androgen receptor, we hypothesised that the most likely target of the repressive effect of testosterone in the pituitary gland is directly on the lactotrophs. We aimed to identify the cell type that androgens act on to repress prolactin release by examining lactotroph-specific, Pit1 lineage-specific and neural-specific conditional AR knockout mice to attempt to duplicate the phenotype of the PARKO. Since the cellular ultrastructure of lactotrophs is sexually dimorphic in rodents (Takahashi and Miyatake, 1991), we aimed to characterise the ultrastructure of the pituitary lactotrophs with electron microscopy to discover whether they develop a characteristic ‘male’ or ‘female’ structure. We also hypothesised that the increase in plasma prolactin in the PARKO may be due to the stimulatory effects on the pituitary of unopposed estrogen signalling, and investigated this by castrating PARKO mice to remove estrogen signalling and quantifying the expression of estrogen stimulated genes in the PARKO pituitary.

## Materials and methods

### Ethics statement

Animal breeding, maintenance and experimental procedures were approved by University of Edinburgh Animal Welfare and Ethical Review Body and were carried out under project license 70/8804 held by Professor Lee B. Smith in line with the UK Home Office Animals (Scientific Procedures) Act, 1986.

### Breeding and maintenance of transgenic mice

Mice in which AR has been ablated selectively from different cell types were generated using Cre/*loxP* technology. In each case, males carrying a Cre transgene were mated to C57BL/6J female mice homozygous for a floxed *Ar* (De Gendt et al., 2004). The *Ar* gene is X-linked. Male offspring were all AR^fl/y^. Those that carried the inherited Cre transgene were defined as conditional knockouts, those that did not were defined as littermate controls.

To generate whole pituitary androgen receptor knockout (PARKO) mice, male congenic 129svev mice carrying a carrying a targeted insertion of Cre recombinase at the F*oxg1* locus (Hebert and McConnell, 2000) were used. To generate lactotroph-specific androgen receptor knockout (Prl-ARKO), mice male mixed of mixed C57BL/6, C57BL/10, and CBA/Ca b with a random insertion of a *Prl*-cre transgene (Castrique et al., 2010) were used. To generate Pit1 lineage-specific androgen receptor knockout (Pit1-ARKO) mice, male mice of a mixed background carrying a carrying a random insertion of *Pou1f1*-cre (Cheung et al., 2018a) were used. Plasma was obtained from neural and glial cell specific androgen receptor knockout mice (Neu-ARKO) used in a previous study (Patel et al., 2017), in which Neu-ARKO mice were generated using male mice of a mixed C57BL/6 and SJF2 background with a random insertion of Nestin-Cre (Tronche et al., 1999). Sex and genotype ratios were all identified at the expected Mendelian ratios. Mice were fed a soya-free diet to avoid any phenotypic effects from dietary estrogens.

### Castration

Mice were anaesthetised by Isoflurane administered via inhalation. A single 1 cm incision was made into the scrotum and testes exposed and removed. Following removal of testes, the site of incision was closed with sterile sutures. Mice were injected subcutaneously with Buprenorphine 0.05mg/kg, whilst anaesthetised, and allowed to recover whilst being monitored. Non-castrated controls were subjected to ‘sham’ surgery where mice were anaesthetised and the incision was made, but testes were not removed. Mice were closely monitored over 24 hours for any welfare problems, and twice daily from then onwards. Mice were culled two weeks after surgery.

### Tissue collection and processing

Brain tissues were fixed for immunohistochemistry following a modified previously published protocol (Rebourcet et al., 2016). Mice were euthanised using a terminal dose of sodium pentobarbital (150 mg/kg, intraperitoneal). Anterograde perfusion fixation of the vasculature was achieved via the left ventricle. First, heparinised PBS (heparin, 20 U/mL) was perfused at 6 mL/min for 2 minutes, then the perfusate was changed to 4% buffered formaldehyde and perfusion continued until blanching of tissues was complete. The brain was then removed and immersion fixed in 4% buffered formaldehyde for another 24 hours, then processed and embedded in paraffin wax. 5μm sections were used for histological analysis.

For all other experiments, mice were euthanised between 14:00h and 17:00h by inhalation of carbon dioxide and subsequent cervical dislocation. Blood was taken from the heart and/or thoracic cavity shortly after cervical dislocation with a needle and syringe coated with heparin, then centrifuged at 20,000 g for 10 minutes at 4°C, then the plasma removed and transferred to storage in a −80°C freezeruntil needed. Forfreezing of pituitary tissue for qRT-PCR, pituitaries were removed and placed on dry ice before being transferred to a −80°C freezer for storage. For fixation of pituitary tissue for electron microscopy, pituitaries were removed and fixed in 2.5% glutaraldehyde and 2% paraformaldehyde in 0.1M phosphate buffer, pH 7.2 at room temperature for 3 hours, then transferred to 0.25% glutaraldehyde and 0.2% paraformaldehyde in 0.1M phosphate buffer, pH 7.2 and then kept at 4°C.

### PCR genotyping of mice

Genomic DNA from ear clips obtained at weaning were subjected to PCR amplification to identify the presence or absence of a Cre transgene. For Foxg1-Cre, Pit1-Cre and Nestin-Cre, the following primers were used: forward: GCGGTCTGGCAGTAAAAACTATC, reverse: AGGCCAGGTATCTCTGACCA, amplicon size: 695 bp). For Prl-Cre mice the following primers were used: forward: CCTGGAAGATGCTCCTGTCTG, reverse: AGGGTGTTGTAGGCAATGCC, amplicon size: 400 bp). Interleukin-2 gene primers were also included in the reaction as a positive control to ensure that DNA was present: Forward: CTAGGCCACAGAATTGAAAGATCT Reverse: AGGCCAGGTATCTCTGACCA, amplicon size: 330bp). PCR was performed with Biomix red kit (Bioline) and PCR products were resolved using the QIAxcel capillary system (QIAGEN, Crawley, United Kingdom).

### Electron microscopy

Pituitaries were prepared for electron microscopy as previously described (Christian et al., 2007). Briefly, cells were post-fixed in osmium tetroxide (1% wt/vol in 0.1 M sodium phosphate buffer), contrasted with uranyl acetate (2% wt/vol in distilled water), dehydrated through increasing concentrations of ethanol (70–100%) and embedded in Spurr’s resin. Ultra-thin sections (50–80 nm) were viewed with a JEM-1010 transmission electron microscope (JEOL USA Inc., Peabody, MA, USA).

### ELISA

Plasma prolactin concentration was measured using a commercially available ELISA (Abcam #ab100736). Plasma prolactin concentrations were expressed as relative change in concentration of conditional knockouts compared to the mean of the controls.

### Preparation of cDNA

Dissected whole pituitaries were frozen on dry ice before being transferred to a −80°C freezer for storage. RNA was isolated using the RNeasy Mini extraction kit (Qiagen, Crawley, UK) according to the manufacturer’s instructions. RNA was quantified using a NanoDrop 1000 spectrophotometer(Thermo Fisher Scientific, Waltham, MA, USA). Random hexamer-primed cDNA was prepared using the SuperScript VILO cDNA synthesis kit (Life Technologies) according to manufacturer’s instructions.

### Quantitative RT-PCR (qPCR)

Multiplex qPCR was performed on pituitary cDNA for the genes listed in Table 1 using an ABI Prism 7900 Sequence Detection System (Applied Biosystems) and the Roche Universal Probe library (Roche, Welwyn, UK). The expression of all genes was related to an internal housekeeping gene assay for *Actb* (Roche, Welwyn, UK) as described previously (O’Hara et al., 2014). Resulting data were analysed using the ΔΔCt method.

**Table 1:**
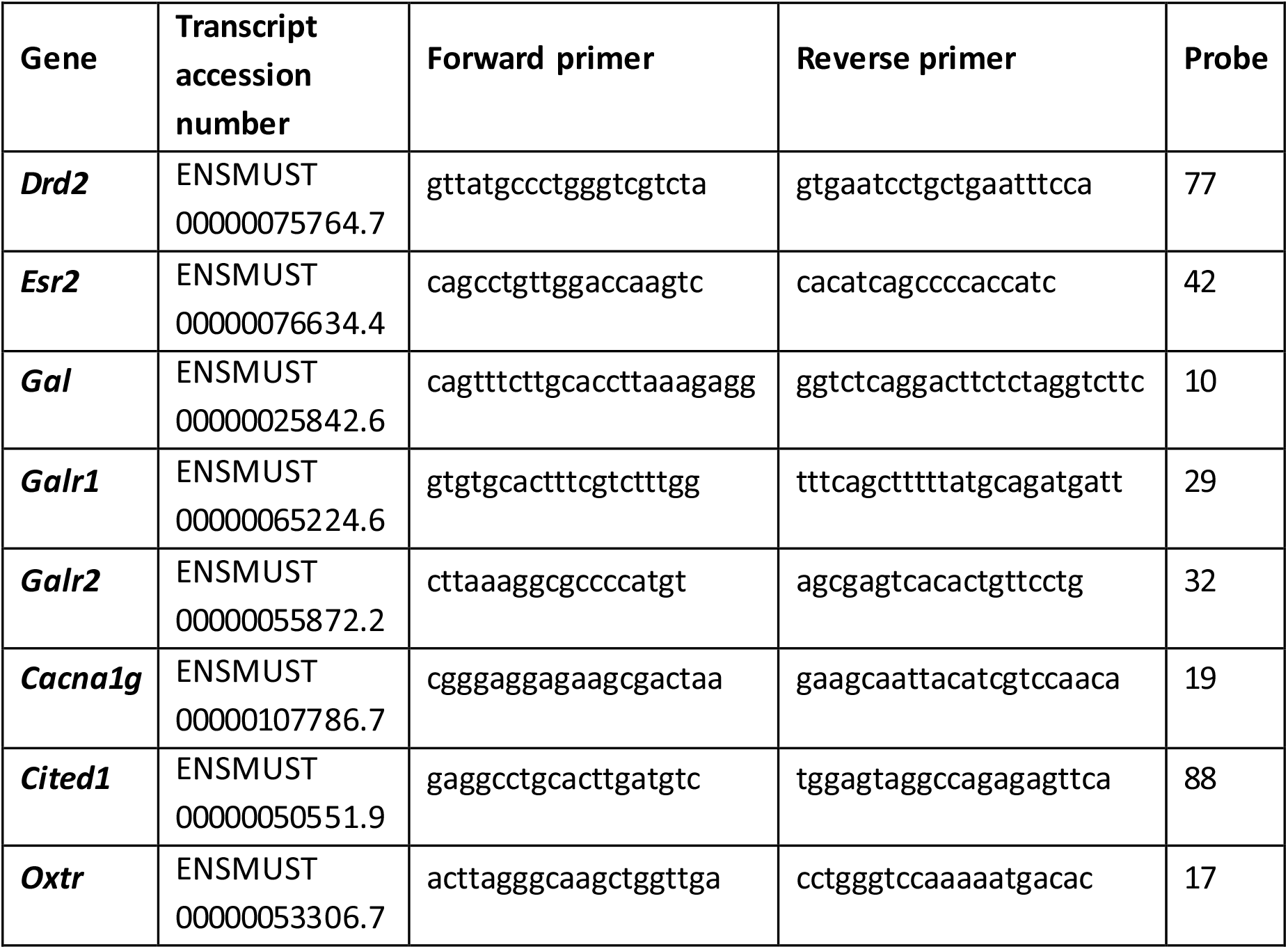
Primers and Roche UPL probes for qRT-PCR assays.

### Immunohistochemistry

Immunolocalisation for androgen receptor and tyrosine hydroxylase (TIDA neurons) was performed by a double antibody tyramide fluorescent immunostaining method, based on a method described previously (O’Hara et al., 2014). Sections of brain prepared as above were deparaffinised and rehydrated, and high-pressure antigen retrieval was performed in 0.01M pH6 citrate buffer for 5 minutes. Non-specific antibody binding sites were blocked with 10% normal goat serum and 1% BSA in TBS (NGS/TBS/BSA). Sections were incubated overnight in a humidity chamber at 4°C in a mixture of rabbit anti-androgen receptor (Spring Bioscience m4070) and chicken anti-tyrosine hydroxylase (Abcam ab76442) in NGS/TBS/BSA. The next morning slides were washed in TBS and endogenous peroxidase was blocked using 0.3% hydrogen peroxide in TBS. The slides were then incubated for 1 hour in a humidity chamber in a mixture of goat anti-rabbit peroxidase conjugated (Vector Labs PI-1000). and goat anti-chicken Alexafluor 488 conjugated (Abcam #ab150169). antibodies both diluted 1 in 200 in NGS/TBS/BSA. Finally the slides were incubated with Cy3 Tyramide Signal Amplification system (‘TSA™’, Perkin Elmer) to manufacturer’s instructions for 10 minutes at room temperature. Slides were mounted with Fluoroshield mounting medium with DAPI (ab104139). Images were captured using a LSM 710 confocal microscope (Zeiss) with Zen software.

### Statistical analyses

Data were analysed using GraphPad Prism (version 7; GraphPad Software Inc., San Diego, CA, USA). If comparing two groups, a two-tailed unpaired t test with Welch’s correction was used for parametric data or a Mann-Whitney test for non-parametric data. If comparing more than one group data were tested for normality and a one-way ANOVA with Bonferroni post-hoc test was used. Graphs display means ± standard deviation error bars.

## Results

### Lactotroph-specific or Pit1 lineage-specific androgen receptor knockout models do not duplicate the increase in circulating prolactin seen in the PARKO

A lactotroph-specific androgen receptor knockout male mouse (Prl-ARKO) was analysed to investigate whether it duplicated the increase in circulating prolactin seen in the PARKO model. Prl-ARKO mice do not have an increase in circulating prolactin compared to control littermates (Figure 1A).

**Figure 1:**
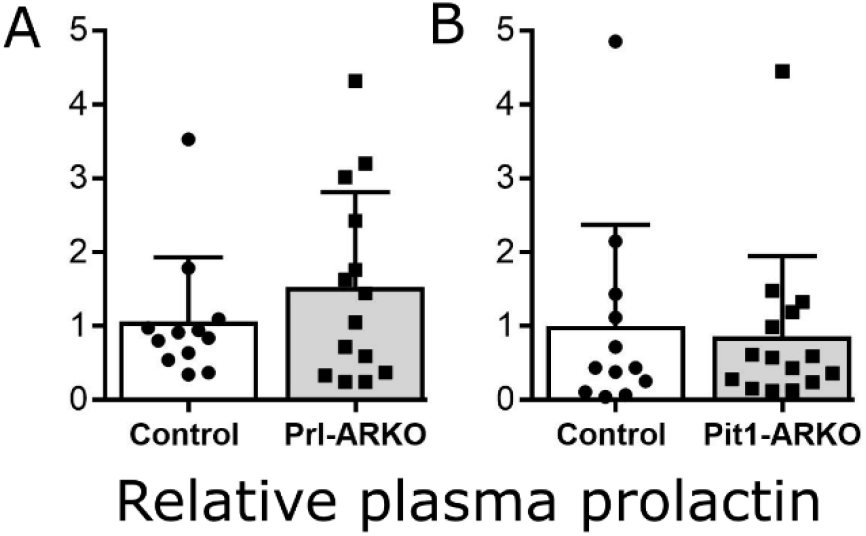
Lactotroph-specific or Pit1 lineage-specific androgen receptor knockout models do not duplicate the increase in circulating prolactin seen in the PARKO. **A:** Prl-ARKO mice do not have a significant change in relative plasma prolactin compared to control littermates (Mann-Whitney test, p>0.05). **B:** Pit1-ARKO mice do not have a significant change in circulating prolactin compared to control littermates (Mann-Whitney test, p>0.05).

Cells expressing the transcription factor Pit1 arise at e13.5 in the pituitary and develop into the lactotroph, somatotroph and thyrotroph populations (Li et al., 1990). A Pit1-Cre line was used to breed a Pit1 lineage-specific receptor knockout male mouse (Pit1-ARKO). This was analysed to investigate whether it duplicated the increase in circulating prolactin seen in the PARKO model. Pit1-ARKO mice also do not have an increase in circulating prolactin compared to control littermates (Figure 1B).

### The increase in circulating prolactin seen in the PARKO is unlikely to result from changes to neural control of prolactin production

Release of prolactin from lactotrophs is under the tonic inhibition of dopamine produced by the TIDA neurons of the arcuate nucleus of the hypothalamus. Although Foxg1-Cre is not generally active in developing hypothalamus, it has been shown to be expressed ectopically in some mouse lines (Hebert and McConnell, 2000). Therefore it was investigated whether an ablation of androgen receptor in the brain was causing TIDA neurons to produce less dopamine and therefore reducing the inhibition on lactotrophs resulting in an increase of prolactin production.

Sections of brains from the region of the hypothalamic arcuate nucleus were double -labelled with both an androgen receptor antibody and a tyrosine hydroxylase antibody to visualis e localisation. TIDA neurons were shown to express androgen receptor in both control (Figure 2A-B) and PARKO (Figure 2C-D) mice. To ascertain whether hypothalamic dopamine content differed between PARKO and control mice, it was measured by ELISA after tissue homogenisation and catecholamine extraction. This showed that there was no change in hypothalamic dopamine between PARKO mice and littermate controls (Figure 2E). Furthermore, plasma from male mice from a neural and glial precursor-cell specific androgen receptor knockout line (Neu-ARKO) was analysed to investigate whether it duplicated the increase in circulating prolactin seen in the PARKO model. Neu-ARKO mice did not have an increase in circulating prolactin compared to control littermates (Figure 2F).

**Figure 2:**
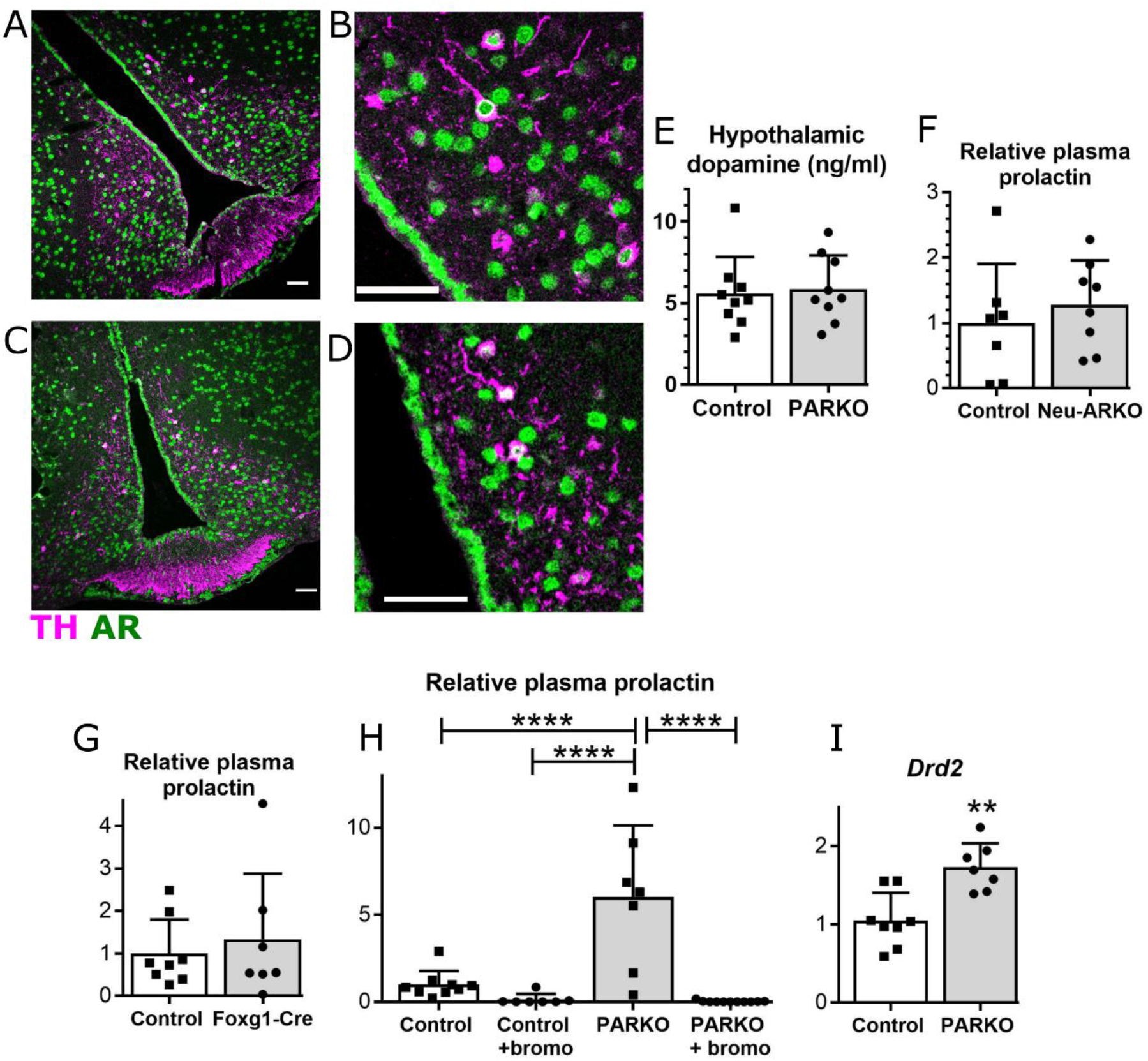
The increase in circulating prolactin seen in the PARKO is unlikely to result from changes to neural control of prolactin production. **A-D:** Sections of brains from the region of the hypothalamic arcuate nucleus were double - labelled with both an androgen receptor antibody (green) and a tyrosine hydroxylase antibody (pink) to visualise localisation. Scale bars 50μm **A:** Control male mouse TIDA neurons express androgen receptor **B:** (Higher magnification of A). **C:** PARKO male mouse TIDA neurons express androgen receptor. **D:** (Higher magnification of C). **E:** PARKO mice do not have a significant change in hypothalamic dopamine compared to control littermates (Mann-Whitney test, p>0.05). **F:** Neu-ARKO mice do not have a significant change in circulating prolactin compared to control littermates (T-test, p>0.05). **G:** Foxg1-Cre mice do not have a significant change in circulating prolactin compared to control littermates (Mann-Whitney test, p>0.05). **H:** Treatment with bromocriptine resulted in a decrease in circulating prolactin in both control and PARKO mice (One-way ANOVA, ****= p<0.0001). **I:** PARKO mice have a significant change in pituitary *Drd2* transcript compared to control littermates (T-test, **= p<0.01).

Foxg1-Cre mice are haploinsufficient as the transgene is a targeted insertion of Cre. This haploinsufficiency has been shown to cause various neural phenotypes, including increases in the hypothalamic neuropeptides oxytocin and arginine vasopressin (Frullanti et al., 2016). To ensure that the increase in plasma prolactin seen was not due to the effects of the Foxg1-Cre by itself, circulating prolactin was measured in Foxg1-Cre mice and compared to littermate controls. However, there was no significant difference between the two groups (Figure 2G).

Finally, to investigate whether the cause of a lactotroph increase in prolactin production in the PARKO model could be tempered by increased agonism of dopamine receptors on the surface of the lactotrophs, mice were given the DRD2 agonist bromocriptine. Treatment with bromocriptine resulted in a decrease in circulating prolactin in both control and PARKO mice (Figure 2H). However, Drd2 transcript was shown to be increased in the PARKO compared to the control (Figure 2I).

### PARKO pituitary lactotrophs in males have a ‘female-like’ lactotroph distribution

Electron microscopy has allowed three types of lactotroph to be characterised in rodents. In rats, type I lactotrophs contain irregularly-shaped granules with diameter 300-700 nm. Type II lactotrophs contain smaller (200-250nm) rounder granules and type III cells contain the smallest (100-200 nm) round granules (Nogami and Yoshimura, 1982). This classification can also be applied in mice (H. C. Christian, unpublished observations). Sexual dimorphism in lactotroph type becomes apparent as rodents age. Both male and female pre-pubertal rat pituitaries contain mostly type III lactotrophs. In early puberty both males and female pituitaries contain mostly type II lactotrophs. By late puberty/early adulthood sexual dimorphism is apparent, with male pituitaries containing approximately 40% type I lactotrophs, 50% type II and 10% type III, compared to females, which contain approximately 80-90% type I, 10-20% type II and 2-4% type III lactotrophs (Takahashi and Miyatake, 1991). To investigate whether the increase in plasma prolactin correlated to any ultrastructural changes, Pituitaries were prepared for electron microscopy and the number of type I lactotrophs (identified with appearance similar to Figure 3 A-B) and type II lactotrophs (identified with appearance similar to Figure 3 C-D) were counted (Figure 3E). Control male mouse pituitaries had on average 50% type I lactotrophs and 50% type II lactotrophs. PARKO male mouse pituitaries had on average 94.25% type I lactotrophs and 4.75% type II lactotrophs.

**Figure 3:**
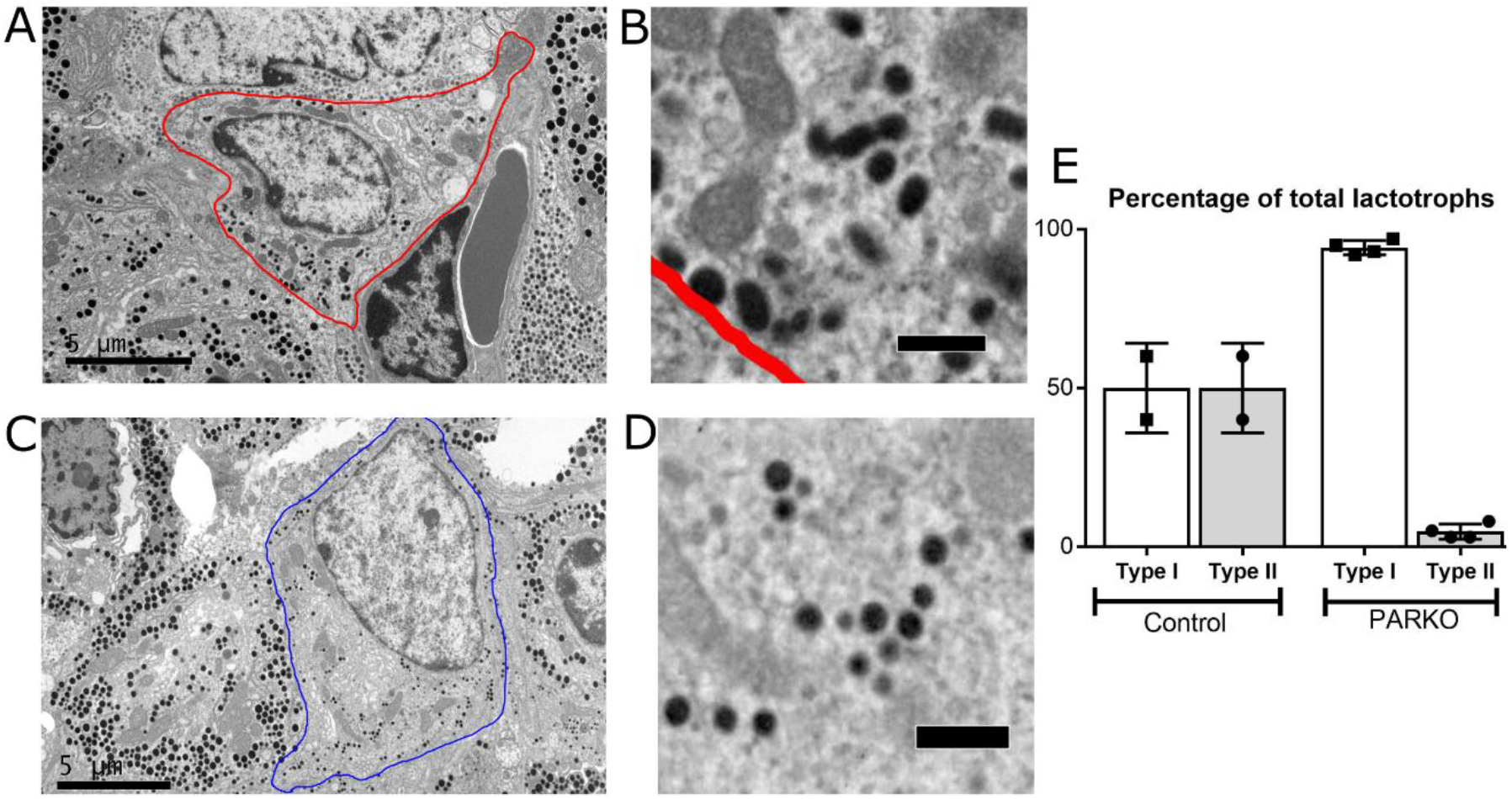
PARKO pituitary lactotrophs in males have a ‘female-like’ lactotroph distribution. **A-D:** Sections of pituitary were visualised by electron microscopy. **A:** Type I lactotrophs can be identified by irregularly-shaped granules with diameter 300-700 nm. Scale bar 5 μm. **B:** Higher magnification of A, scale bar 0.5 μm. **C:** Type II lactotrophs can be identified by smaller (200-250nm) rounder granules. Scale bar 5 μm. **D:** Higher magnification of C, scale bar 0.5 μm. **E:** Control male mouse pituitaries had on average 50% type I lactotrophs and 50% type II lactotrophs. PARKO male mouse pituitaries had on average 94.25% type I lactotrophs and 4.75% type II lactotrophs.

### Castration reduces circulating prolactin in PARKO mice to that of castrated controls, but not intact controls

Pituitaries from male rats treated with estrogen have majority type I lactotrophs, unlike non estrogen-treated rats that have a majority type II (De Paul et al., 1997). Since this is similar to PARKO male pituitaries we hypothesised that the higher circulating prolactin in the PARKO mouse was due to unopposed estrogen signalling in the pituitary. To investigate whether the balance between androgen and estrogen signalling is important in pituitary prolactin release, control and PARKO mice were castrated, allowed to recover for two weeks and then their circulating prolactin was compared to sham operated littermates (Figure 4A). Castration increased circulating prolactin in control mice by 4.4x (p>0.05, not significant). However, this was not as high as intact PARKO mice which had mean circulating prolactin 8.6x higher than that of uncastrated control mice (*** p<0.001), and this significantly dropped to 3.432x control after castration (** p<0.01).

**Figure 4:**
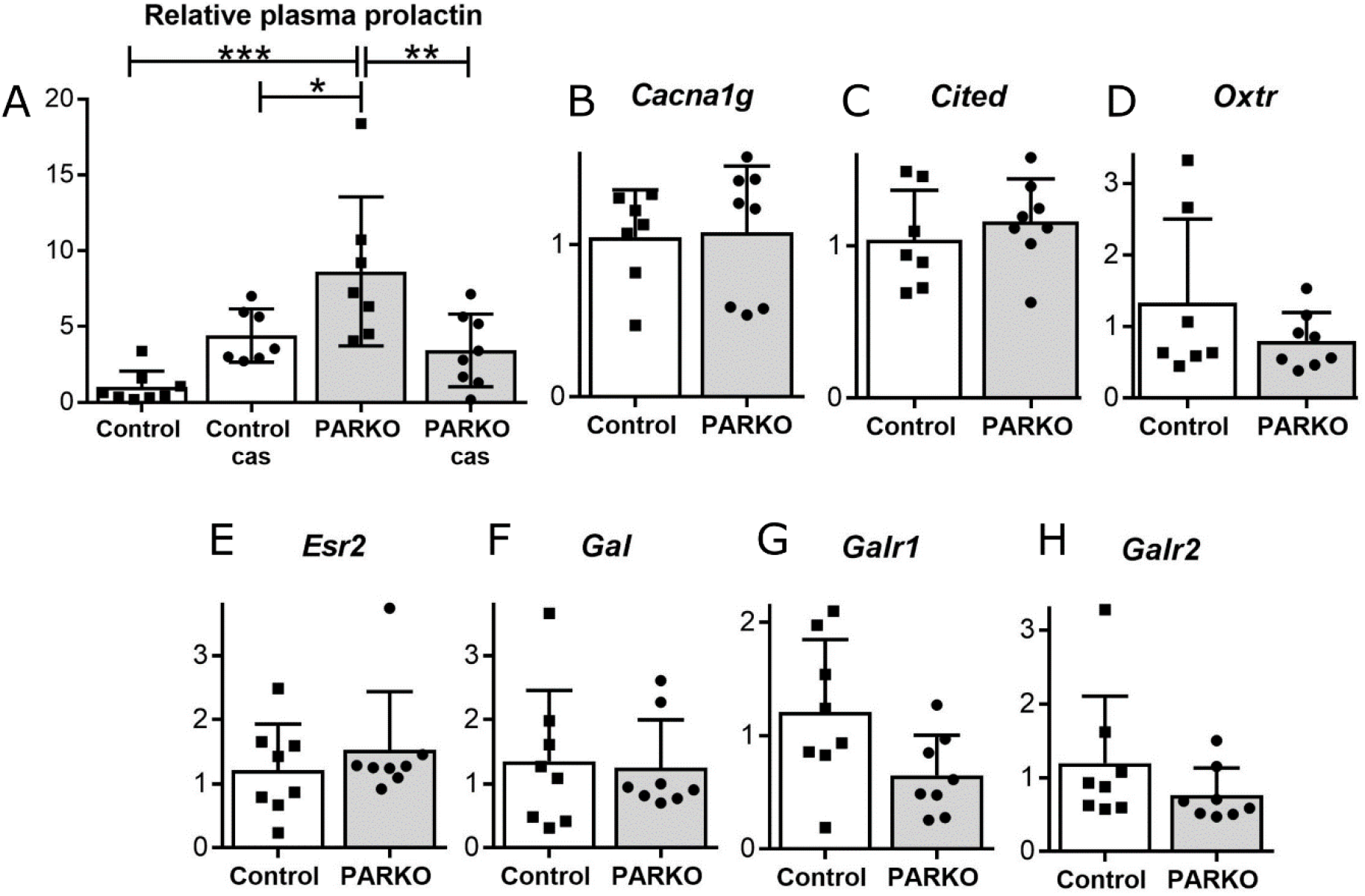
Castration reduces circulating prolactin in PARKO mice to that of castrated controls, but not intact controls. A: Castration did not significantly increase circulating prolactin in control male mice (p>0.05, not significant). Uncastrated PARKO mice had mean circulating prolactin 8.6x higher than that of uncastrated control mice (*** p<0.001), and this significantly dropped to 3.432x control after castration (** p<0.01). B: Calcium channel, voltage-dependent, Ttype, alpha 1G subunit (*Cacna1g*) expression was not significantly different in PARKO pituitaries compared to controls (Mann-Whitney, p>0.05). C: Cbp/p300-interacting transactivator with Glu/Asp-rich carboxy-terminal domain 1 (*Cited1)* expression was not significantly different in PARKO pituitaries compared to controls (T-test, p>0.05). D: Oxytocin receptor (*Oxtr*) expression was not significantly different in PARKO pituitaries compared to controls (Mann-Whitney, p>0.05). E: Estrogen receptorbeta (*Esr2*) expression was not significantly different in PARKO pituitaries compared to controls (Mann-Whitney, p>0.05). F: Galanin (*Gal* expression was not significantly different in PARKO pituitaries compared to controls (T-test, p>0.05). G: Galanin receptor 1 *Galr1* expression was not significantly different in PARKO pituitaries compared to controls (T-test, p>0.05). H: Galanin receptor 2 *Galr2* expression was not significantly different in PARKO pituitaries compared to controls (Mann-Whitney, p>0.05).

*Esr1* expression (estrogen receptor alpha) was previously shown to be unchanged in the PARKO (O’Hara et al., 2015). However, this does not show if signalling through ERα is changed. Calcium channel, voltage-dependent, T type, alpha 1G subunit (*Cacna1g*), Cbp/p300-interacting transactivator with Glu/Asp-rich carboxy-terminal domain 1 (*Cited1)* and oxytocin receptor (*Oxtr*) are three genes previously identified to be upregulated by estradiol/ ERα in the mouse pituitary (Kim et al., 2011). However the expression of these three genes was not different in PARKO compared to control (Figure 4B-D), suggesting that signalling through ERα has not been increased.

Estrogen signalling can also be transmitted through *Esr2* (estrogen receptor beta). *Esr2* expression was also shown to be unchanged in PARKO mice compared to controls (Figure 4E). The expression of the components of the galanin pathway were also investigated. Galanin is a neuropeptide secreted by the pituitary upregulated by estrogen to stimulate the release of prolactin (Kaplan et al., 1988). Expression of Galanin (*Gal*, Figure 4F) and two of its receptors *Galr1* (Figure 4G) and *Galr2* (Figure 4H) were not significantly different between control and PARKO mice.

## Discussion

We have previously identified androgens as a novel inhibitor of prolactin release by characterisation of a mouse line with ablation of androgen receptor in the pituitary during fetal development (pituitary androgen receptor knockout: ‘PARKO’) (O’Hara et al., 2015). This resulted in an increase in circulating prolactin concentration in male mice, despite no increase in lactotroph number. In this study we have shown that PARKO male mice develop lactotrophs that resemble those seen in female mice, and that this is likely to contribute to the increase in circulating prolactin. The role of androgens as a inhibitor of prolactin release is under-investigated in the literature, whereas estrogen is a well-established stimulator of prolactin release. In one study, DHT has previously been shown to reverse the stimulatory effect of estradiol on *Prl* mRNA levels in rats. It is yet to be determined whether this repression is directly through androgen receptor binding to the prolactin promoter or through an indirect mechanism. Testosterone treatment has been known to supress lactation in post-partum women since the 1930s (Kurzrok and O’Connell, 1938). This was shown to be at least in part at the level of hypothalamic-pituitary control rather than direct effects on the breast, as serum prolactin levels decreased with testosterone treatment (Weinstein et al., 1976). Another human study has shown that treatment of trans women with the androgen receptor antagonist cyproterone acetate is responsible for their increase in plasma prolactin before orchiectomy is performed (Defreyne et al., 2017).

Electron microscopy has facilitated the characterisation of three types of lactotroph in rodents. Type I lactotrophs contain irregularly-shaped granules with diameter 300-700 nm. Type II lactotrophs contain smaller (200-250nm) rounder granules and type III cells contain the smallest (100-200 nm) round granules (Nogami and Yoshimura, 1982). In adulthood, sexual dimorphism of lactotroph type is apparent, with male pituitaries containing approximately 40% type I lactotrophs, 50% type II and 10% type III, compared to females, which contain approximately 80-90% type I, 10-20% type II and 2-4% type III lactotrophs. This sexual dimorphism is thought to be generated by ovarian estrogen production at puberty. When adult male rats are treated with estrogen they develop a female-specific lactotroph distribution, and when adult female rats are ovariectomised they develop a male -specific distribution (Takahashi and Miyatake, 1991).

In the PARKO model, we are not changing the levels of circulating sex steroids but rather removing any androgen signalling at the pituitary by ablating androgen receptor expression. This causes males to produce and release more prolactin and for the lactotrophs to develop a female-specific distribution. We hypothesised that, although circulating estrogen has not changed, removal of androgen signalling in the pituitary would have caused the balance of androgen-estrogen signalling in the pituitary to be pushed more towards the estrogen-dominant ‘female’ state. Although estrogen production in PARKO males would be low, its action at the pituitary would be unopposed by any counteracting androgen signalling. We hypothesised that removal of circulating estrogens and androgens (as estrogen precursors) by castration would redress this balance. When castrated, PARKO males have significantly reduced circulating prolactin compared to intact males. However, we did not investigate lactotroph distribution in castrated PARKO mice and it would be interesting to discover whether the mechanism of the circulating prolactin reduction was a change of lactotroph distribution from a more ‘female-specific’ pattern back to a more ‘male-specific’ one.

To investigate further the balance of estrogen signalling in these mice, we looked at the expression of three genes that have been previously characterised as upregulated by estradiol/ERα in the mouse pituitary, namely: calcium channel, voltage-dependent, T type, alpha 1G subunit (*Cacna1g*), Cbp/p300-interacting transactivator with Glu/Asp-rich carboxy-terminal domain 1 (*Cited1)* and oxytocin receptor (*Oxtr*) (Kim et al., 2011). However the expression of these three genes was not different in PARKO compared to control, suggesting that signalling through ERα has not been increased. This is perhaps unsurprising: even if signalling through estrogen receptor has not been increased, it could still be greater than any counteracting effect provided by androgen receptor signalling. We also looked at the underlying physiological mechanism of the prolactin increase in PARKO males. Galanin is a neuropeptide secreted by the pituitary that is upregulated by estrogen to stimulate the release of prolactin (Kaplan et al., 1988). Expression of Galanin and two of its receptors *Galr1* (Figure 4G) and *Galr2* were investigated but were not shown to be significantly different between control and PARKO mice suggesting that this system is not contributing to the prolactin increase in PARKO males. The recent advent of pituitary single-cell RNA sequencing studies has provided a wealth of information about the transcriptome differences between rodent pituitary cell types (Ho et al., 2020, Cheung et al., 2018b), and also sexual dimorphism in pituitary gene expression (Fletcher et al., 2019). The Fletcher et al. study showed that all endocrine cell types and folliculostellate cells in rats have genes that are dominantly expressed by sex, with lactotrophs having the most at 288. Careful analysis of the data generated from these sequencing studies may help future elucidation of both the mechanisms of sexualdimorphism of pituitary prolactin production, and the role of androgen receptor in its control.

The PARKO model is a general ablation of androgen receptor in the pituitary gland. Since lactotrophs express androgen receptor and produce prolactin, we investigated whether a targeted ablation of androgen receptor specifically in the lactotrophs could duplicate the phenotype seen in the PARKO. This was first explored in a ‘Prl-ARKO’ mouse model, in which androgen receptor was ablated by a Cre driven a 3.2 kb fragment of the rat PRL promoter. There was no increase in circulating prolactin in the Prl-ARKO mice compared to littermate controls. The Prl-Cre has been shown to be more efficient in female mice, where it targets 70-80% of lactotrophs (Fu and Vankelecom, 2012, Castrique et al., 2010), than in male mice, where it targets 35% of lactotrophs (Fu and Vankelecom, 2012). Since this incomplete targeting may have been preventing the duplication of the phenotype, we switched to using a Pit1-Cre to ablate androgen receptor. Cells expressing the transcription factor Pit1 arise at e13.5 in the pituitary and develop into the lactotroph, somatotroph and thyrotroph populations (Li et al., 1990). Despite this earlier, more complete ablation of the androgen receptor, we could not duplicate the increase in circulating prolactin concentration seen in the complete PARKO mouse model.

It is also possible that the increase in circulating prolactin seen in the PARKO has been caused by ablation of androgen receptor by Foxg1-Cre in the brain. The most likely neural cell type that would affect prolactin production and release are the TIDA neurons of the hypothalamus, that produce inhibitory dopamine. Although the Foxg1-Cre used to produce the PARKO model is not generally active in developing hypothalamus, it has been shown to be expressed ectopically in some mouse lines (Hebert and McConnell, 2000). Therefore we investigated whether an ablation of androgen receptor in the brain was causing TIDA neurons to produce less dopamine and therefore reducing the inhibition on lactotrophs resulting in an increase of prolactin production. Several pieces of evidence generated from this study lead us to conclude that the TIDA neurons are not targeted by Foxg1-Cre, and that hypothalamic dopamine release is not affected by androgen receptor ablation. Although TIDA neurons express androgen receptor, there is no ablation in PARKO mice and PARKO mice do not have an increase in hypothalamic dopamine. Also, mice with a complete ablation of androgen receptor in neural and glial cells (Neu-ARKO) do not have an increase in circulating prolactin. This evidence suggests that it may not be androgen receptor signalling in the lactotroph that is negatively regulating prolactin production, but in another cell type in the pituitary. Since androgen receptor in the somatotrophs and thyrotrophs will also be ablated with the Pit1-Cre, this leaves either the gonadotrophs, corticotrophs or folliculostellate cells as likely candidates. Paracrine signalling between gonadotrophs and lactotrophs is well characterised (Denef, 2008), but signalling between corticotrophs and lactotrophs is less well-characterised. There is also the possibility that the increase in circulating prolactin seen in the PARKO is not attributable to just one cell type, and is the result of the interaction of changes to several cell types in the pituitary when androgen receptor signalling during pituitary development is disrupted. The final possible explanation is that the increase in plasma prolactin seen in the PARKO is due to an off-target knockout of androgen receptor in a tissue other than the pituitary or hypothalamus.

Further investigations are needed into the mechanism of prolactin regulation by changes in androgen-estrogen balance, which has implications not only in the normal sexual dimorphism of physiology but also in diseases such as hyperprolactinemia. Pathological hyperprolactinemia can have a numberof causes, including pituitary prolactinoma. However, in males it is often associated with low circulating testosterone levels (De Rosa et al., 2003). Although this can be caused by prolactin repression of GnRH neuron activity, the association between cause and effect is not fully characterised. Defining the interaction of prolactin with the HPG axis will be necessary for fully understanding reproductive function and disorders.

## Acknowledgements

### Funding

This work was supported by the Biotechnology and Biological Sciences Research Council (grant number BB/N007026/1).

## Acknowledgements

We thank Dr. Saloni Patel for collection of plasma from Neu-ARKO mice, Dr. Diane Rebourcet for assistance with perfusion fixation of mice, Dr. Nicola Romanò for advice on brain immunostaining, Jyoti Nanda, Dr. Laura Milne and Nathan Jeffrey for laboratory technical assistance, Michael Dodds for animal husbandry support and Lynne Scott for assistance with preparing sections for electron microscopy.

